# Tracking stem cell differentiation without biomarkers using pattern recognition and phase contrast imaging

**DOI:** 10.1101/097329

**Authors:** John D. Delaney, Yuhki Nakatake, D. Mark Eckley, Nikita V. Orlov, Christopher E. Coletta, Chris Chen, Minoru S. Ko, Ilya G. Goldberg

## Abstract

Bio-image informatics is the systematic application of image analysis algorithms to large image datasets to provide an objective method for accurately and consistently scoring image data. Within this field, pattern recognition (PR) is a form of supervised machine learning where the computer identifies relevant patterns in groups (classes) of images after being trained on examples. Rather than segmentation, image-specific algorithms or adjustable parameter sets, PR relies on extracting a common set of image descriptors (features) from the entire image to determine similarities and differences between image classes.

Gross morphology can be the only available description of biological systems prior to their molecular characterization, but these descriptions can be subjective and qualitative. In principle, generalized PR can provide an objective and quantitative characterization of gross morphology, thus providing a means of computationally defining morphological biomarkers. In this study, we investigated the potential of a pattern recognition approach to a problem traditionally addressed using genetic or biochemical biomarkers. Often these molecular biomarkers are unavailable for investigating biological processes that are not well characterized, such as the initial steps of stem cell differentiation.

Here we use a general contrast technique combined with generalized PR software to detect subtle differences in cellular morphology present in early differentiation events in murine embryonic stem cells (mESC) induced to differentiate by the overexpression of selected transcription factors. Without the use of reporters, or *a priori* knowledge of the relevant morphological characteristics, we identified the earliest differentiation event (3 days), reproducibly distinguished eight morphological trajectories, and correlated morphological trajectories of 40 mESC clones with previous micro-array data. Interestingly, the six transcription factors that caused the greatest morphological divergence from an ESC-like state were previously shown by expression profiling to have the greatest influence on the expression of downstream genes.

## Introduction

Since the development of the compound microscope in the late 1500s, characterizing morphological complexity has been the central goal of many biologists. In the interceding five centuries modern microscopy has advanced to the point where it is dependent on high-powered computers for image acquisition, data storage and analysis. As a result of this technological convergence, “big data” experiments involving large scale image processing have become the prevailing approach to biological pathway discovery where over-expression, knockdown, GFP-tagging, drug screens, etc., all rely heavily on high throughput systems designed to capture and analyze unprecedented quantities of image data.

Researchers interested in studying poorly characterized biological processes are often challenged with the lack of appropriate biomarkers. Commonly used ionic dyes and lysochromes (e.g. Haematoxylin/Eosin, Oil Red O, etc) can be of great benefit when initiating characterization of a new system but consistent staining can be difficult to achieve for very large numbers of samples. Alternatively, phase contrast or differential-interference contrast (DIC) imaging has the advantage of not relying on specific contrast agents, uniform staining or extensive sample preparation, and can rapidly provide uniform data sets ideal for large-scale experiments.

While acquisition of these image sets is straightforward, automated analysis can be quite complicated. Segmentation algorithms have been used for analyzing phase contrast and DIC imaging, and some recent examples include automated mitosis detection (Huh et al. 2011) and tracking neural stem cells (Winter et al. 2011). However, segmentation depends entirely on the accuracy of detecting the objects of interest, which relies on manual or iterative fine-tuning of algorithm parameters. Although used successfully in the past (Cohen et al. 2010), this type of image analysis strategy can be especially problematic when used to identify individual cells in the heterogeneous populations characteristic of cell differentiation studies. The high developmental cost associated with robust segmentation renders scalability a major issue.

The combination of generic contrast techniques with general purpose non-parametric image processing can facilitate the routine use of imaging assays in large scale experiments, without requiring expertise in developing or refining image-specific analysis algorithms, and without the need for specific molecular markers for the processes being studied (Tarca et al. 2007), (Peng 2008), (Logan & Carpenter 2010), (Shamir, et al. 2010).

The application of machine learning and pattern recognition to image data, originally developed for remote sensing, has been increasingly used for biological image processing (Boland et al. 1998) (Shariff et al. 2010), (Brown et al. 2002), (Neumann et al. 2006), (Collinet et al. 2010), (Eliceiri et al. 2012). We have developed a PR program (WND-CHARM) with a specific emphasis on generality of input image types while minimizing external parameter adjustments required from the user (N. Orlov et al. 2008), (Shamir, et al. 2008). The key image parameters specified to WND-CHARM are limited to assigning input images into groups (classes), e.g., defining experimental controls, standard curves, time courses, etc.

WND-CHARM can discern complex patterns in gross morphology without *a-priori* knowledge of what those patterns are; allowing analysis of images generated with non-specific contrast methods (DIC, phase-contrast). WND-CHARM also reports quantitative measures of similarity (i.e. distance) between image classes or individual images, allowing it to be used in quantitative assays where qualitative class assignment is inadequate. Some of the studies where WND-CHARM has been previously used include: identification of aging states in *C. elegans* using DIC (Johnston et al. 2008), (Eckley et al. 2013), classification of lymphoma types (N. V. Orlov et al. 2010a) and characterization of melanoma progression (N. V. Orlov et al. 2012) using H&E-stained sections.

The study of stem cell differentiation into several different lineages imaged by phase contrast can pose an extreme challenge to existing automated image processing techniques. In this study, we use phase contrast imaging and PR-based analysis to track the earliest morphological changes in 40 different transgenic murine embryonic stem cell clones (mESC) by imaging them over a time course. Each mESC clone carries a different inducible transcription factor selected based on its importance in early embryonic development (Nishiyama et al. 2009). Induction of these different transcription factors drives these mESC clones along different differentiation trajectories (Aiba et al. 2009). The phenotypes detected by the software (representing the fate of each clone) are consistent with the predicted cell fates based on previous expression profiling studies of these cell lines. Additional DIC imaging of the cells in the collection as well as their molecular characterization can be found online at the: NIA Mouse ES Cell Bank (http://esbank.nia.nih.gov/publications.html). The data sets used in the work presented here can be found online at the Image Informatics and Computational Biology Unit (IICBU) website: (http://ome.grc.nia.nih.gov/indexworking.html).

While we have previously reported the use of this software for classifying images of cells, the unique contribution of this study is that we now show that quantitative comparison of phenotype similarity using general pattern recognition techniques can be used to distinguish and characterize multiple morphological trajectories without the use of molecular biomarkers. The characterization of these trajectories using gross morphology alone was consistent with more detailed molecular studies. Furthermore, neither the analysis software nor its parameters was specific to the cells or imaging modality used in this study, and the contrast method used here can be applied to any experimental system. This study is a demonstration that PR and its objective interpretation of gross morphology can be used to characterize biological processes where conventional biomarkers are unavailable. Moreover, although this software is internally more complex than conventional segmentation-based approaches, its reliance on example images for user input rather than parameter and algorithm selection can make it easier for experimental biologists to use.

## Results

### Morphological progression of murine embryonic stem cells

Initially we sought to quantify the divergence of a transcription factor (TF) induced mESC clone away from an Ntg (empty vector) parental control. The cells were grown for seven days in standard embryonic stem cell media supplemented with Leukemia inhibitory factor (LIF) and the presence (gene-off) or absence (gene-on) of Doxycycline (Dox). In this series of experiments images were captured daily on the same cell cultures. To gauge the effect of inducing each gene over time compared to the Ntg control, we used pair-wise vector subtraction (see Materials and Methods for explanation of method) to calculate the Morphological Divergence Score (MDS) between each gene at each time point and its corresponding time point for the Ntg control. Thus the MDS reports a divergence from Ntg as a result of overexpressing the chosen gene while accounting for morphological differences due solely to Dox or genetic manipulation.

The results from these experiments are represented graphically in figure 1. Although each cell line diverges from the Ntg, the rate and degree of progression differs for each gene. As a negative control we calculated the MDS between Nanog and Pou5f1 (Oct4), two clones that have been shown to signal through the same pathway to maintain pluripotency in ESC (Liang et al. 2008) and therefore should have very similar phenotypes. We found that these two clones were identical to the Ntg parental cells and each other for the first three days of the experiment and showed very little progression away from each other in days five and seven (MDS of 0.027 and 0.021 respectively). This shows that a low MDS results after overexpression of genes that are not expected to diverge from Ntg or from each other. The MDS between Nanog and Oct4 at day seven (0.021) constitutes a cutoff for significant morphological differences due to differentiation. By these criteria the earliest detectable significant morphological change occurs at day three for Gata3.

**Figure1:**
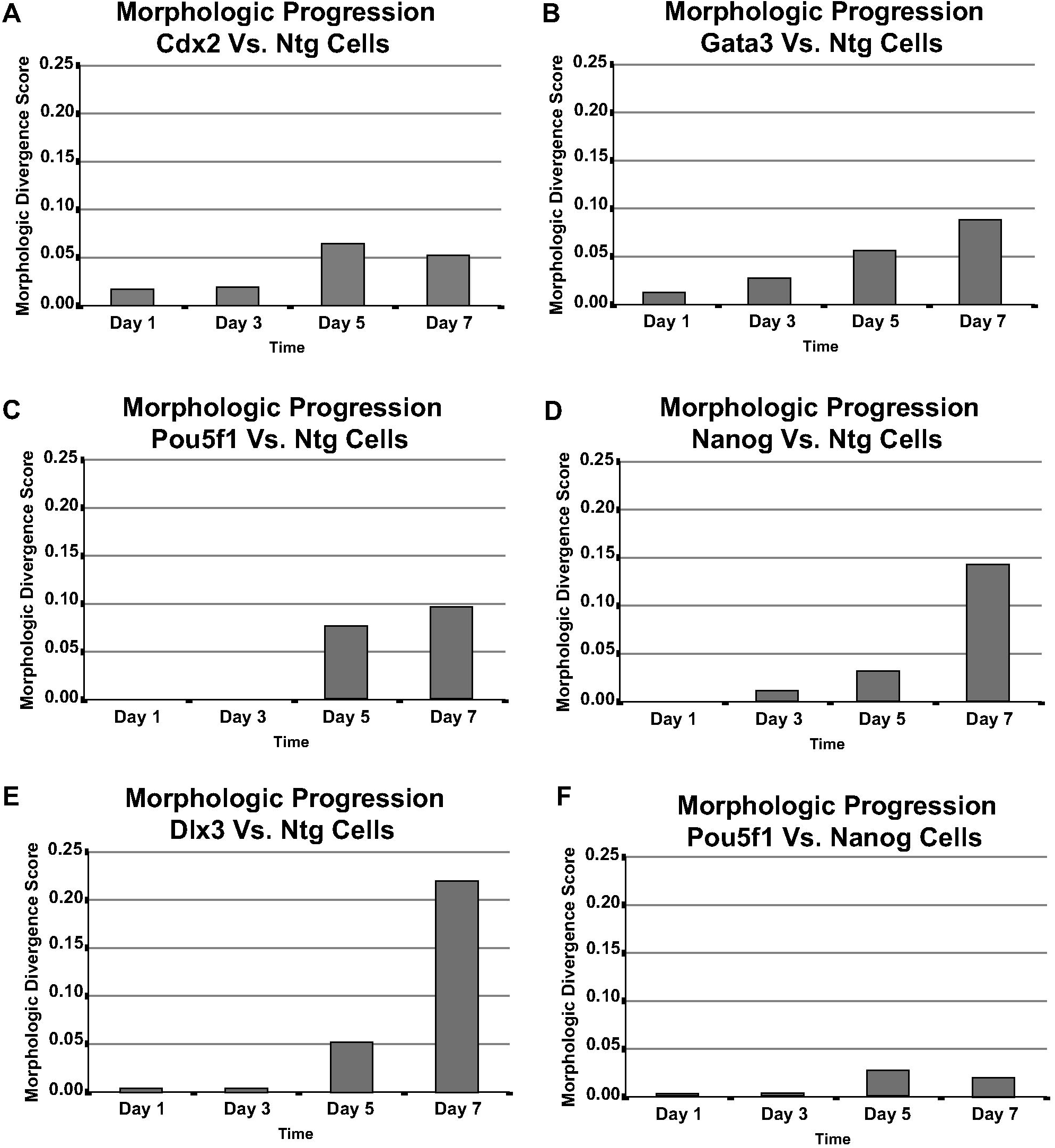
Morphologic divergence of transgenic murine embryonic stem cells from an Ntg control. Distances between the induced clones and the Ntg controls are measured as a MDS (see methods). 1A: Progression of Cdx2. 1B: Progression of Gata3. 1C: Progression of Pou5f1/Oct4. 1D: Progression of Nanog. 1E: Progression of Dlx3. 1F: Control experiment, divergence of Nanog from Pou5f1/Oct4. Nanog and Oct4 are known to function through the same pathway to maintain pluripotency.

### Phylogenetic representation of eight mESC clones after seven days of TF induction

Having quantified mESC differentiation over time, we tested the reproducibility of WND-CHARM by comparing three similar experiments performed independently. In these experiments all the clones were grown for seven days and images were captured at the termination of the experiment (representative images of clones tested in supplemental figure 1). One of the variables to be considered when growing eight different cells lines under two different culture conditions is differences in rate of cell growth. Although these cells lines are very similar, activation of the transgene can affect proliferation rates to varying extents depending on the gene expressed. To address this issue we curated the images (as discussed in Materials and Methods) to restrict the gene comparisons to image samples (tiles) with comparable cell density.

Initially 400 image samples were collected of each condition (i.e. 25 fields of view), but after pre-classification screening, the sample number for each condition ranged from 125 to 398. In order to not bias the classifier, we limited the number of training images for each class, as determined by the class with the smallest number of images. To visualize the morphological similarities between the classes, we used the program PHYLIP to create phylogenic trees based on the pair-wise distances between classes reported by WND-CHARM. Figure 2A represents data from the first biological replicate. The limiting class in this experiment contained only 125 images so all classes used 100 images for training and 25 images for testing. Figure 2B is the second biological replicate, here 185 images were used for training and 25 for testing. Figure 2C is the third biological replicate with the greatest number of images, 299 were used for training and 25 for testing. As there were a large number of images to consider and WND-CHARM uses a random selection for populating the training and testing sets, each experiment was performed 1000 times, and the final figures are based on the average distances from these trials.

**Figure 2:**
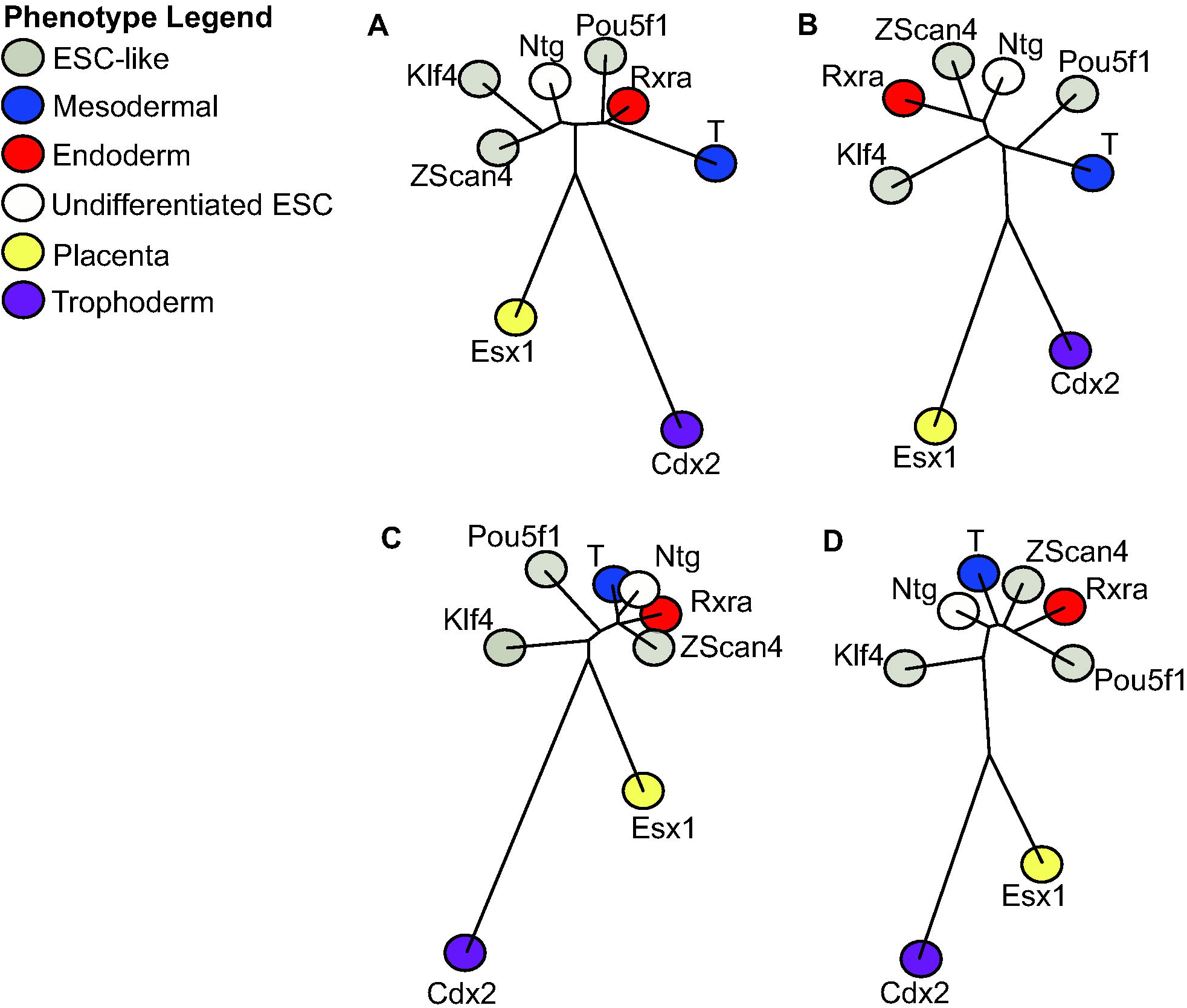
Visualization of distances calculated by WND-CHRAM between murine embryonic stem cell clones. The dendrograms represented here are visual representations of WND-CHARM distance calculations for three replicate eight-gene experiments (2a, 2b and 2c). The high reproducibility between experiments is demonstrated by the similar relative positions of the genes across experiments. Significantly, both Cdx2 and Esx1 segregate away from the central body of gene phenotypes while maintaining their proximity to each other. Both of these genes are implicated in placenta and trophectoderm development, which is the first divergence of cell fates in a developing embryo. Figure 2d was constructed from all three replicates using 125 images per class. The genes are identified in these dendrograms by the addition of color circles at the terminus of branches indicating different cell fates based on previously established functional associations.

While there was a modest amount of shuffling of some of the genes in the three replicates (Zscan4, Pou5f1, Rxra) there was also a high degree of reproducibility. In all three of the replicates both Cdx2 and Esx1 map away from the main body of phenotypes. Interestingly both Cdx2 and Esx1 are important in trophectoderm / placental development. In fact trophectoderm is one of the first fate switches that occurs in a developing mass of embryonic cells. Here, three independent classifiers reproducibly segregated out two cell populations, based solely on a gross morphological phenotype, that have similar genetic roles / functions. Next, a composite analysis was performed where 125 image samples from each class in each replicate were pooled across the replicates, generating classes with 375 samples. These pooled samples were then used to create a fourth independent classifier with 350 training images and 25 testing images. Once again the experiment was performed 1000 times and the results plotted with PHYLIP, Figure 2D.

It is difficult at this stage to definitively identify subsequent fates, such as mesoderm or endoderm. This is not particularly surprising, as this test only represents seven transcription factors, which may not represent sufficient morphological divergence. To address this, we expanded our experiment to a larger set of transcription factors.

### Phylogenetic Representation of 40 mESC clones after seven days of TF induction

Reproducibly classifying phenotypes into the same relative positions in a multi-dimensional space with four distinct classifiers indicated that it would be possible to combine many experiments into a master dendrogram.

Figure 3 is a compilation dendrogram using newly acquired image data comparing phenotype similarity among 39 cell lines expressing the indicated TF and an Ntg control. Even though the classifier only had phase contrast images to work with, it was able to show that cells expressing genes with similar functional associations have similar morphologies, and are therefore closely spaced in the dendrogram. For example, N-Myc, c-Myc and Smad4 form a tight cluster in the dendrogram in Figure 3. Functionally, although both N-Myc and c-Myc genes are essential for completion of murine embryonic development, the two proteins are highly homologous and replacement of c-myc coding sequences with those from N-Myc can lead to normal development, indicating that the two loci facilitate differential patterns of expression of these functionally similar proteins (Malynn et al. 2000). Smad4 has been demonstrated to bind and activate c-myc promoter via the positive regulatory element TBE1. Thus the observed morphological cluster in Figure 3 can simply be a functional consequence of three alternative mechanisms of increasing myc expression (Lim & Hoffmann 2006).

**Figure3:**
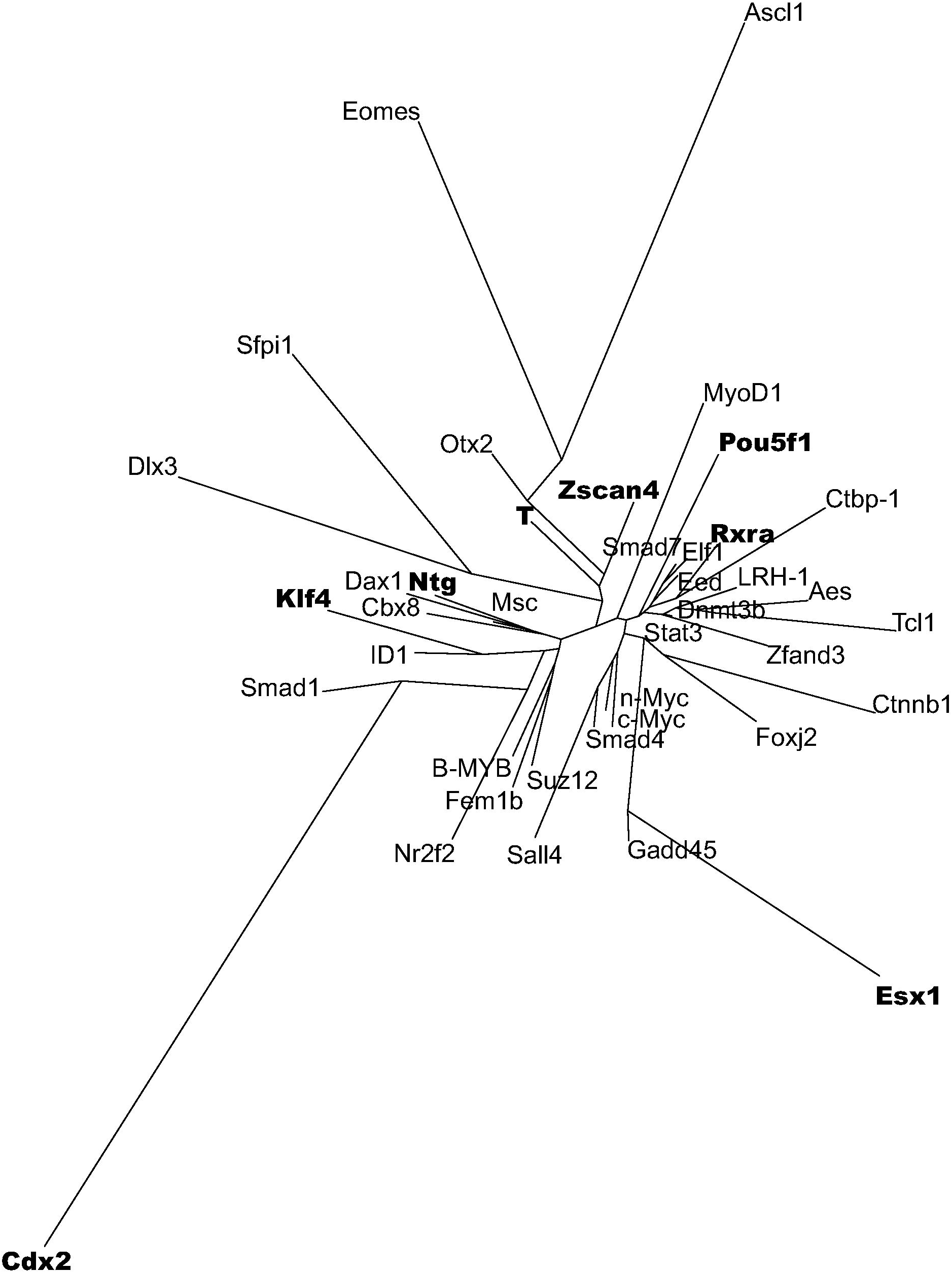
Visualization of morphologic distances calculated by WND-CHARM for 40 murine embryonic stem cell clones. In this experiment, six clones (Ascl1, Cdx2, Dlx3, Eomes, Esx1 and Sfpi1) were able to escape the ESC-like cloud in the center of the dendrogram. In previous work (Nishiyama et al. 2009) these same clones were able to affect the transcription of the greatest number of downstream genes.

The relative positions of the cell lines used in the previous eight-gene experiment are also reproduced here with newly acquired images, and it can be seen that their relative placement is preserved and not affected by the additional 32 clones. In this figure the previous clones (Cdx2, Esx1, Klf4, Ntg, Pou5f1, Rxra, T and Zscan4) are bolded.

## Discussion

The genes explored in this study were chosen from an initial microarray analysis of Oct3/4 signaling (Matoba et al. 2006), as well as gene regulatory network profiling (Aiba et al. 2009) performed on the same cell lines as those used in this study. The results we present here based on morphologic data alone are consistent with previous results based on transcription profiles of these cell lines (Aiba et al. 2009). Additionally, principle component analysis (Nishiyama et al. 2009) showed that the transcriptome state for the TF-induced ESCs is stable for the first 48 hours. Similarly, in figure 1 we show that the earliest observable differences in morphology induced by TF overexpression occurs at day three.

The results in Figure 3 demonstrate it is possible to push an ESC out of a pluripotent state by overexpressing individual TFs even in the presence of LIF, which is normally used to inhibit differentiation in ESC culture. Examples of the TFs that can cause this escape phenotype include Cdx2 and Esx1 (trophectoderm), Dlx3 and Ascl1 (ectoderm), Eomes (endoderm) and Sfpi1 (mesoderm). The previous work (Nishiyama et al. 2009) analyzing transcription profiles of these cell lines measured the number of affected genes as an indicator of TF potency. The six TFs we identified that were able to escape the ESC-like state were shown in the previous work to affect the transcription of the greatest number of genes, indicating that TF potency is well correlated with the ability to drive morphologic differentiation. The one obvious difference between the two studies is the effect of overexpressing Klf4. In the previous work, Klf4 affected the expression of over 2500 genes, but in our study it was not able to induce the escape phenotype. The major difference between Klf4 and the six TFs identified here is that KLF4 is already abundantly expressed in ESCs, so while additional levels of Klf4 may indeed further perturb the expression levels of many downstream genes, this may not result in a differentiation phenotype. Furthermore, the ESC-like phenotype of Klf4 overexpression is consistent with it’s role together with Oct4, Sox2 and c-Myc in reprograming somatic cells into pluripotent stem cells (Takahashi & Yamanaka 2006).

A persistent problem in many biological systems, exemplified in differentiating ESCs, is that experimental perturbations result in heterogeneous cultures. Analyzing heterogeneous cultures in bulk (by e.g. microarrays, proteomic analysis, etc.) results in average characteristics for the culture, which in the worst case, may not apply to any of its constituent cells. The analysis presented here shows that a machine can be trained to identify several different differentiation pathways at the same time. Here, we studied differentiation in multiple cultures, but the same approach can be used to examine differentiation within a single culture, after being trained on several separate cultures. This technique can be used analytically to investigate the nature of the heterogeneity (e.g. co-expression of various biomarkers), or it can be combined with cell isolation techniques (e.g. an image-based cell sorter) to obtain collections of homogenous cells for subsequent molecular characterization.

Quantitative image analysis is a core technology in biological investigation. A major impediment to its wider adoption is the effort and expertise required to design imaging algorithms and tune their parameters to each type of cell or imaging subject, imaging modality, contrast method, and experimental perturbation. An alternative is to use non-parametric pattern recognition algorithms to quantify differences between experimental subjects while accounting for variation within experimentally defined samples. Here, the sole parameter is the arrangement of images into treatment groups, and the machine automatically identifies differences between groups while ignoring differences within groups. Any imaging technique and subject can be used in the analysis. The ability to quantify morphological differences between experimental groups regardless of the imaging methodology and without the need to develop new algorithms or adjust parameters can be the basis for a much more general approach to image analysis, and a much wider adoption of imaging technology in routine biological assays.

The image data and experimental system used in this study was chosen to represent an extreme case of heterogeneity within experimental samples together with a contrast technique that does not rely on any previously-known biomarkers for use as specific probes. The difficulty of this test case was further compounded by the need to distinguish and track differences along several different morphological trajectories within the same experiment. Many biological assays are in fact simplifications of what is presented here. For example, figure 1 presents a time course of differentiation along several morphological trajectories at the same time, each induced by a different transcription factor. A more common assay is a time-course of a single perturbation, or equivalently, a set of concentrations in a dose-response experiment of a single drug. Images in a simpler assay can be grouped by time point or drug dose to obtain a quantitative response curve based on morphological similarity. In previous work, we have applied this software to a variety of imaging problems, modalities and subjects. These range from brightfield and fluorescence microscopy (Shamir, et al. 2008) to histology (N. V. Orlov et al. 2010b), including non-biological problems such as face recognition (N. Orlov et al. 2008). Considering that the only effort required to determine if a particular imaging problem can be addressed using this non-parametric approach is to collect a representative set of training images, we believe that this software can become a routine tool in the modern biological laboratory.

## Material and Methods

### Embryonic Stem Cell Clones

These experiments were performed with clonal populations of the murine embryonic stem cell line MC-1 harboring transcription factors under the control of a ROSA-Tet system (Nishiyama et al. 2009). Cells were grown in a standard embryonic stem cell media; High Glucose (4.5 g/l D-Glucose) DMEM, 15% FBS; LIF (ESGRO, Chemicon, USA) 1000 U/ml; 1 mM sodium pyruvate; 0.1 mM NEAA, 2 mM glutamate, 0.1 mM beta-mercaptoethanol, and penicillin/streptomycin (50 U/50 μg per ml). Transcription factors were expressed by the absence of Doxycycline (0.01ug/ml) in the media. Cells were plated at 1000 cells / well in 96 well plates coated with 0.1% gelatin and grown for up to seven days in standard ESC media ± Doxycycline (Dox). Media was changed every other day. After seven days the cultures were fixed with 4% paraformaldehyde at 4°C over night and then washed with 3x with PBS. Plates were imaged on a GE Incell Analyzer 1000 using the phase-contrast setting at 10x magnification. All images were saved as .tif files.

### Pattern Recognition and the WND-CHARM Software

WND-CHARM relies on extracting and comparing image features to distinguish between two or more classes of images. The software initially computes more than 3000 features from each image and uses a Fisher discriminant to rank order the most informative features and assign them weights. The weighted feature vectors for each image sample are then classified by a modified nearest neighbor classification scheme. Once an image classifier has been trained, the groups of images can be mapped into a multi-dimensional feature space using similarity measurements, where each axis represents one of the user-defined image classes. The resulting data output is either .html or .txt format. (N. Orlov et al. 2008), (Shamir, et al. 2008).

### Generation of Signature Files

All data (.tifs and meta files) was exported from the InCell Analyzer to a centralized server system for analysis. An annotation file was generated from the meta-data that contained all of the gene names and experimental conditions for each plate. In the first phase of the analysis a Perl program (wndchrm_experiment) processed an experimental description file to create and populate a directory structure of images for a given classification problem. For instance, in a two-gene classification problem the program would create directories for gene1+Dox, gene1-Dox, gene2+Dox, gene2-Dox; the WND-CHARM program then processed this directory structure and produced classification results. In these experiments the following parameters were used with WND-CHARM (release 1.31-255): -l (large feature set), -S500 (normalizes the mean pixel intensity to 500), -t4 (tiles the images 4 x 4). These parameters resulted in 16 independently calculated sets of image features for each field of view in each class (one for each of the 16 tiles).

### Controlling for Non-uniform Growth Rates

It was anticipated that the clones would not grow at a uniform rate. To address this issue, images were selected from all classes and manually assigned to a group of confluent cells or bare plastic. Using these two classes a simple two-way classifier was generated. Using the WND-CHARM classify command, all image-feature files representing tiles of the original field of view, were scored by this classifier as either confluent or plastic, with all samples scored as “plastic” removed from further analysis. The resulting curated classes were compared to each other using the WND-CHARM test command. Because each clone exists in an induced and quiescent state (Dox- and Dox+ respectively) the analysis was performed in a pairwise fashion for all genes. For each pair-wise comparison between genes, a four-way classifier was generated, for example: gene1-on, gene1-off, gene2-on, gene2-off to compare gene1 and gene2.

### Morphological Divergence Score (MDS)

After WND-CHARM calculated the marginal probabilities for all four classes in each pair-wise gene comparison, a Perl program (pairwise_distance) was used to process the WND-CHARM HTML reports and collect all marginal probabilities for distance calculations. Each pair-wise gene comparison is done using four classes (gene1-on, gene1-off, gene2-on, gene2-off), which results in four average marginal probabilities for each class, where the averages are computed from 1000 individual runs. The marginal probabilities for each of the four classes are used as coordinates in a four-dimensional space, with one dimension per class. Distances between any two classes are computed as the Euclidean distance in this space. The Morphological Divergence Score between gene1 and gene2 is the result of the gene-off distance (gene1-off to gene2-off) subtracted from the gene-on distance (gene1-on to gene2-on). In this way, only the effect of the transgene being overexpressed is taken into account when constructing a dendrogram of morphologic distances. This process is repeated for each pairwise comparison in the analysis and the results are saved in a final pair-wise distance file. The FITCH and DRAWTREE programs from the PHYLIP (Felsenstein 1989) package were then used to process the pair-wise distance files and generate the dendrograms.

### Algorithm

WND-CHARM is written in C++ and makes use of the public available fftw and libtiff libraries (Frigo & Johnson 2005). The code is compiled using the standard *gcc* compiler and is easily ported to most commonly used operating systems. The WND-CHARM program as well as the Perl utilities described in this paper are available as GPL-licensed free software on Google Code: (http://code.google.com/p/wnd-charm/). WND-CHARM 1.31-255 was used for the analysis of this data.

## Acknowledgements

This research was supported entirely by the Intramural Research Program of the NIH, National Institute on Aging.

## Supplementary Material

**Supplemental Figure 1:**
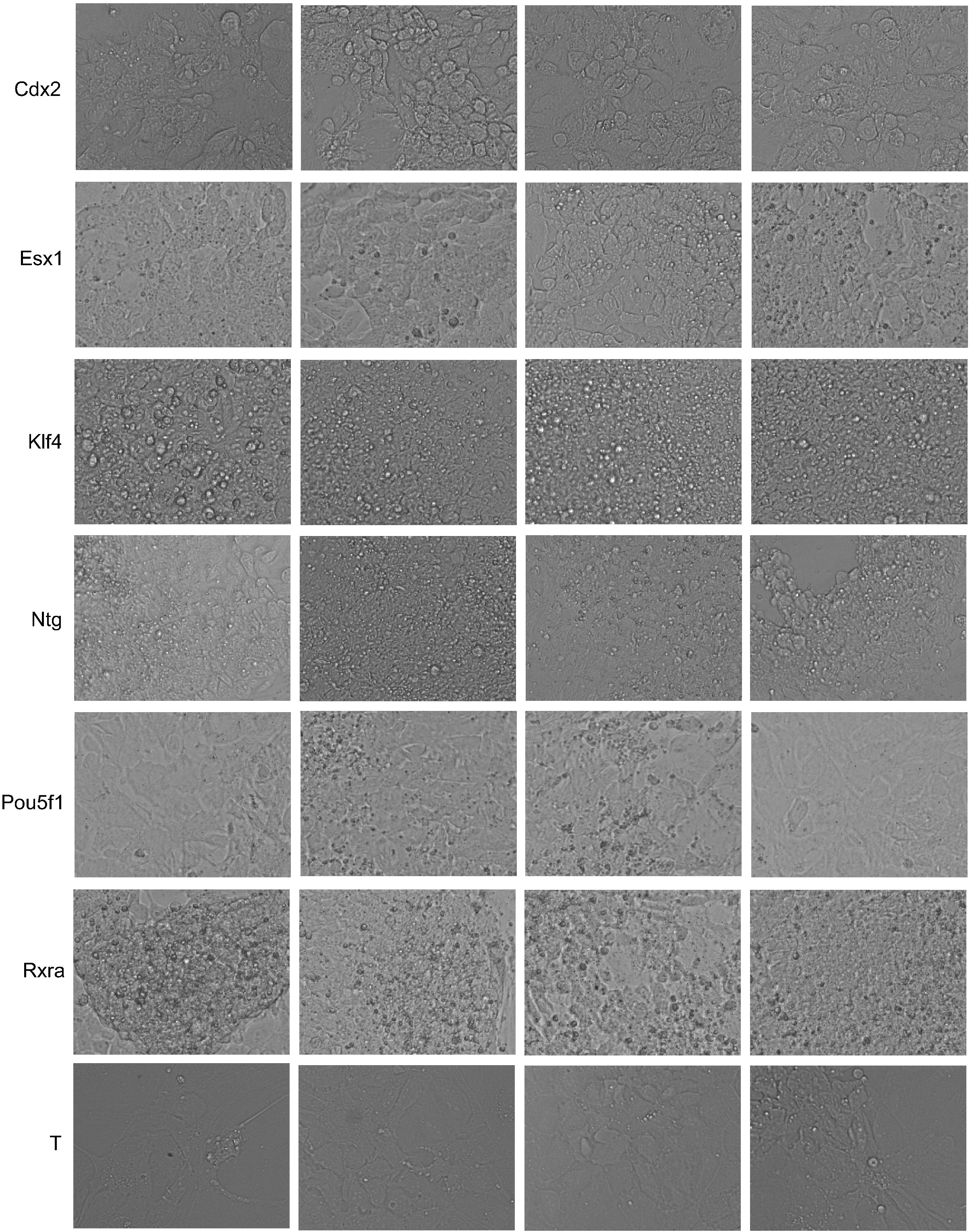
Example morphologies for the transgenic embryonic stem cells used in these experiments. All images are taken from Dox - (induced) stem cell clones at a magnification of 10x. Each box represents one image that was used in the analysis.

